# *Legionella* protection and vaccination mediated by Mucosal Associated Invariant T (MAIT) cells

**DOI:** 10.1101/231472

**Authors:** Huimeng Wang, Criselle D’Souza, Xin Yi Lim, Lyudmila Kostenko, Troi J Pediongco, Sidonia BG Eckle, Bronwyn S Meehan, Nancy Wang, Shihan Li, Ligong Liu, Jeffrey YW Mak, David P Fairlie, Yoichiro Iwakura, Jennifer M Gunnersen, Andrew W Stent, Jamie Rossjohn, Glen P Westall, Lars Kjer-Nielsen, Richard A Strugnell, James McCluskey, Alexandra J Corbett, Timothy SC Hinks, Zhenjun Chen

## Abstract

Mucosal associated invariant T (MAIT) cells recognize conserved microbial metabolites from riboflavin synthesis. Striking evolutionary conservation and pulmonary abundance implicate them in antibacterial host defense, yet their roles in protection against clinically significant pathogens are unknown. Murine *Legionella* infection induced MR1-dependent MAIT cell activation and rapid pulmonary accumulation of MAIT cells associated with immune protection detectable in fully immunocompetent host animals. MAIT cell protection was more evident in mice lacking CD4+ cells, whilst profoundly immunodeficient RAG2^−/−^γC^−/−^ mice were substantially rescued from uniformly lethal *Legionella* infection by adoptively-transferred MAIT cells. This protection was dependent on MR1, IFN-γ and GM-CSF, but not IL-17, TNF-α or perforin. Protection was enhanced and observed earlier post-infection in mice that were Ag-primed to boost MAIT cells before infection. Our findings define a significant role for MAIT cells in protection against a major human pathogen and indicate a potential role for vaccination to enhance MAIT cell immunity.

---

Mucosal-associated invariant T (MAIT) cells are innate-like lymphocytes with the potential to recognize a broad range of microbial pathogens. MAIT cells express a ‘semiinvariant’ αβ T cell receptor (TCR) and recognize small molecules presented by the major histocompatibility complex (MHC) class I-related molecule (MR1)^1, 2^. These molecules comprise derivatives of the riboflavin biosynthetic pathway^3-5^, which is conserved between a wide variety of bacteria, mycobacteria and yeasts^3, 6^, but is absent from mammals, and therefore provides an elegant mechanism to discriminate host and pathogen. Indeed the enzymatic pathway required for riboflavin synthesis has been identified in all microbes shown to activate MAIT cells, and is absent in those that do not^3^.

A striking feature of MAIT cell immunity is the high level of conservation of MR1 across 150 million years of mammalian evolution^7-9^, implying a strong evolutionary pressure to maintain the MAIT cell compartment. Furthermore, MAIT cells have a strong pro-inflammatory phenotype^10^ and are abundant in humans in blood and lung tissue^11^, whilst in C57BL/6 mice are found in greater abundance in the lungs than any other organs^12^. Together these features implicate MAIT cells in a critical role in respiratory host defense. However, very few pathogens have been demonstrated *in vivo* to cause activation and proliferation of MAIT cells^13, 14^. In studies implicating a role for MAIT cells in protective immunity against pathogens, the definition of these cells was limited by the lack of MR1-Ag tetramers^14^. To date no studies have clearly defined a functional role for MAIT cells in protection against a clinically important human pathogen.

Using a model of bacterial lung infection with the intracellular bacteria *Salmonella enterica* serovar Typhimurium we have previously shown that riboflavin gene-competent bacteria can cause rapid activation and proliferation of MAIT cells^13^. We therefore hypothesized that this response could also be elicited with an authentic human lung pathogen and would contribute to protection against disease.

*Legionella spp* are facultative intracellular pathogens, gram-negative, flagellated bacteria which, when inhaled, cause a spectrum of disease from self-limiting Pontiac fever to severe, necrotic pneumonia: Legionnaire’s disease^15^. Incidence of Legionnaire’s has nearly trebled since 2000, with >5000 cases/year in the USA, inflicting a 10% mortality despite best treatment^16^. In North America and Europe^16^ the predominant pathogen is *L. pneumophila* whilst in Australasia and Thailand over 50% of cases are caused by *L. longbeachae^17^*.

Here we have used MR1 tetramers loaded with the potent MAIT cell ligand 5-(2-oxopropylideneamino)-6-D-ribitylaminouracil (5-OP-RU)^18^ to specifically identify^4^ and characterize MAIT cells in human *in vitro* and murine *in vivo* models of lung infection with the two most clinically significant *Legionella* species: *L. pneumophila* and *L. longbeachae.* Our data reveal that MAIT cells contribute to protection against fatal infection with *Legionella*, by a mechanism that is dependent on MR1 and interferon (IFN)-γ / granulocyte macrophage-colony stimulating factor (GM-CSF). Protection is partial in immunocompetent hosts but becomes increasingly evident as other arms of immunity are disabled such as in CD4 T cell-deficient animals. Protection ultimately becomes “all or nothing” in profoundly immunodeficient mice RAG2^−/−^γC^−/−^ mice. These studies dissect the mechanisms by which MAIT cells contribute to protection against an important human disease and a model intracellular pathogen.

## Results

### Human MAIT cells are activated by *Legionella* infection *in vitro* via MR1

We^3, 13^ have previously shown that MAIT cells are activated by microbial species that express the riboflavin biosynthetic pathway; a finding which has been confirmed by others^6^. We therefore investigated whether *Legionella* species known to cause serious pulmonary infections in humans - *L. pneumophila^15^’^19^* and *L. longbeachae^17^* - and to express the necessary *rib* enzymes^20^, could activate human MAIT cells. First, bacterial lysates of *L. pneumophila* and *L. longbeachae* stimulated a reporter cell line expressing a MAIT TCR (Jurkat.MAIT-A-F7)^3^ in the presence of an MR1-expressing lymphoid cell line (C1R.MR1)(Figure 1A). Stimulation was dose-dependent, and could be specifically blocked by anti-MR1 antibody^21^. Next we used a well-characterized human monocytic cell line (THP-1)^22^ as an antigen presenting cell co-cultured with flow-sorted human peripheral blood CD3+Va7.2+CD161+ cells. We observed activation of MAIT cells when co-cultured with THP1 cells infected for 27 hours with live *L. longbeachae* (Figure 1B,C). Intracellular infection of wild type THP1 and THP-1:MR1+ cell lines induced expression of TNF-a by human MR1-5-OP-RU tetramer+ MAIT cells. Activation was related to the infective dose, and was specific to MAIT cells and not non-MAIT CD3+ T cells. Activation was MR1-dependent, as it did not occur in the presence of cells in which we had disrupted the MR1 gene using a CRISPR/Cas9 lentiviral system (THP1:MR1-). MAIT cells also expressed IFN-g in the presence of MR1-over-expressing cells (THP1:MR1+), but expression was minimal using the parental cell line (THP1), which has very low constitutive surface expression of MR1.

**Figure 1.**
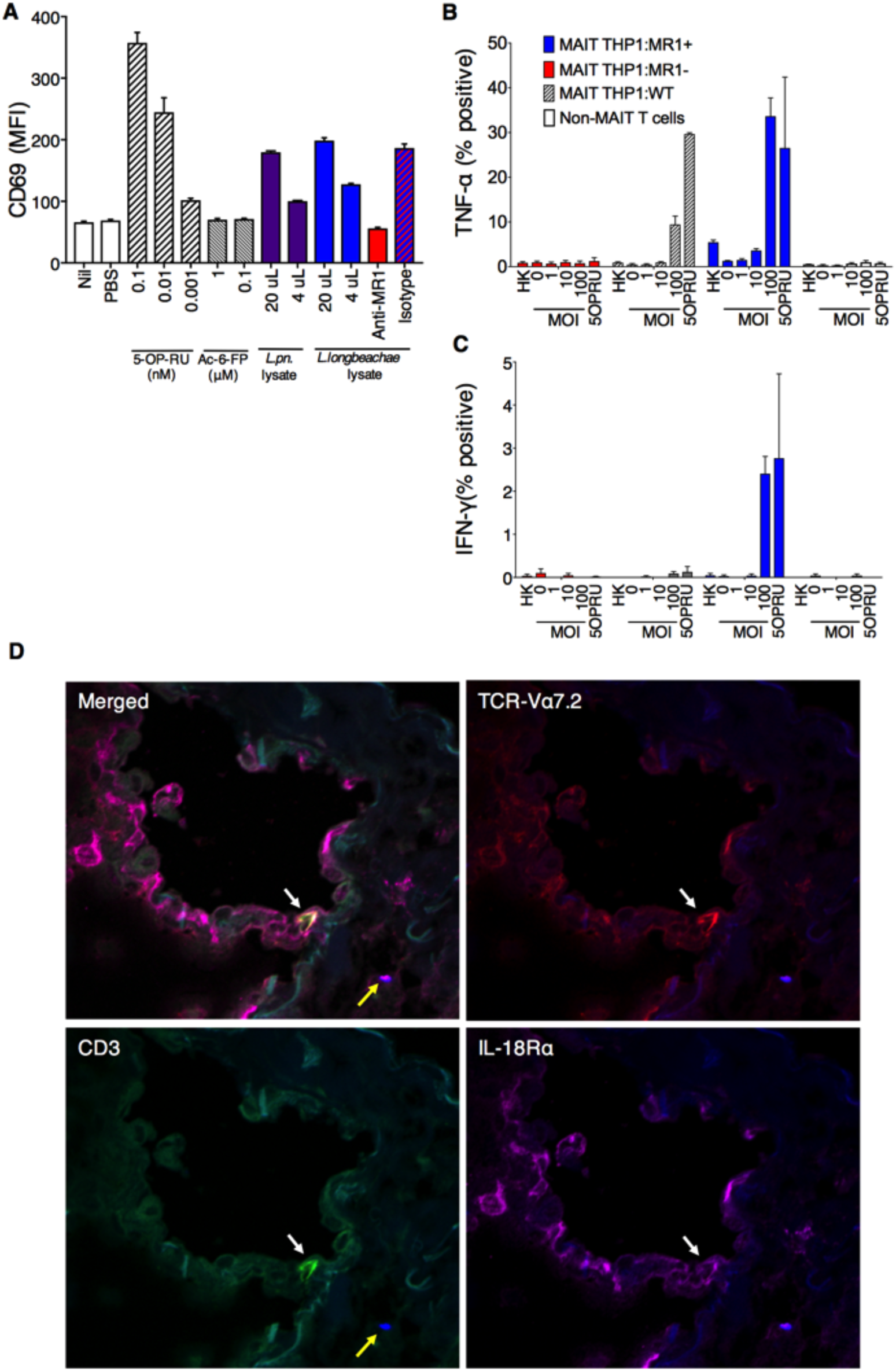
Human MAIT cells are activated by *Legionella* infection via MR1 *in vitro* (A) Jurkat.MAIT and C1R.MR1 cells were co-incubated for 16h with lysates of *L. pneumophila (L. pn.) or L. longbeachae*, or 5-OP-RU, acetyl-6-FP or PBS. Activation, detected by staining with anti-CD69, is enhanced by bacterial lysate or by the activating ligand 5-OP-RU, but not by acetyl-6-FP. Activation was blocked by anti-MR1 antibody (26.5) but not by isotype control (W6/32) 2h prior to co-incubation. (B-C) THP1 cells (WT) or THP1 cells overexpressing MR1 (THP1:MR1+, blue) or deficient in expression of MR1 (THP1:MR1-, red) were infected for 27h with *L. longbeachae*, heat killed (HK) *L. longbeachae* (MOI100) or 10nM 5-OP-RU, then co-cultured for 16h with CD3^+^Va7.2^+^CD161^+^ human peripheral blood MAIT cells, or MAIT-depleted conventional T cells. MR1-5-OP-RU-tetramer+ MAIT cell activation was measured by intracellular cytokine staining for (B) TNF-a or (C) IFN-g. (D) Immunofluorescence micrographs showing CD3+TCRVa7.2+IL-18Ra+ MAIT cell (white arrow) within healthy human lung tissue 24h post infection *ex vivo* with *L. longbeachae* (yellow arrow). Red, TCR-Va7.2; green, CD3; magenta, IL-18Ra; blue, legionella. Data show MFI or percentage cytokine-positive cells with SEM.

To visualize MAIT cells *in situ* we infected healthy human lung tissue *ex vivo* with *L. longbeachae* and observed CD3+TCRVa7.2+IL-18Ra+ MAIT cells within the lung parenchyma in the proximity of *Legionella* bacilli 24 hours post-infection using immunofluorescence microscopy (Figure 1D).

These findings indicate that *Legionella* induces potent MAIT cell immune responses *in vitro* suggesting that MAIT cells are likely to play a role in protection against *Legionella* pneumonia.

### MAIT accumulate in the lungs during *Legionella* infection *in vivo*

Next we examined the impact of *Legionella* infection on MAIT cells *in vivo* in a murine model using intranasal infection with live *L. longbeachae.* TCRß+ MR1-5-OP-RU tetramer+ cells were visible in the lung parenchyma using immunofluorescence microscopy within three days post-infection (Figure 2A). There was striking enrichment of pulmonary MAIT cells (from here on defined as CD45+TCRβ+ MR1-5-OP-RU tetramer+ cells), which comprised up to 30% of all pulmonary αβ-T cells after 7 days (Figure 2B, C). MAIT cell accumulation was dependent on size of initial inoculum and was proportionately much larger for MAIT cells - 580-fold absolute increase at 10^5^ colony-forming units (CFU) (P<0.0001) - than conventional αβ-T cells (maximum 9.4- fold, P<0.0001) (Figure 2C,D). Accumulation occurred rapidly over 7 days post infection (DPI), with absolute numbers peaking at day 10 (Figure 2E). Furthermore, despite a subsequent 20-fold contraction from peak frequencies (P=0.005), overall expansion of the MAIT cell population was long-lived, persisting >280 DPI (Figure 2E,F). Interestingly, although MAIT cells have been implicated in recruitment of non MAIT T cells ^14^, we did not observe any significant difference in pulmonary recruitment of αβ-T cells in MR1-/- mice, which have an absolute deficiency of MAIT cells^12, 13^. Likewise, i.n. infection with 2×10^7^ CFU *L. pneumophila* similarly induced a rapid expansion of MAIT cells (Supplementary Figure S1), although more modest than *L. longbeachae.* As C57BL/6 mice are susceptible to *L. longbeachae^17^, L. longbeachae* was selected as the most appropriate model for more detailed investigation.

**Figure 2.**
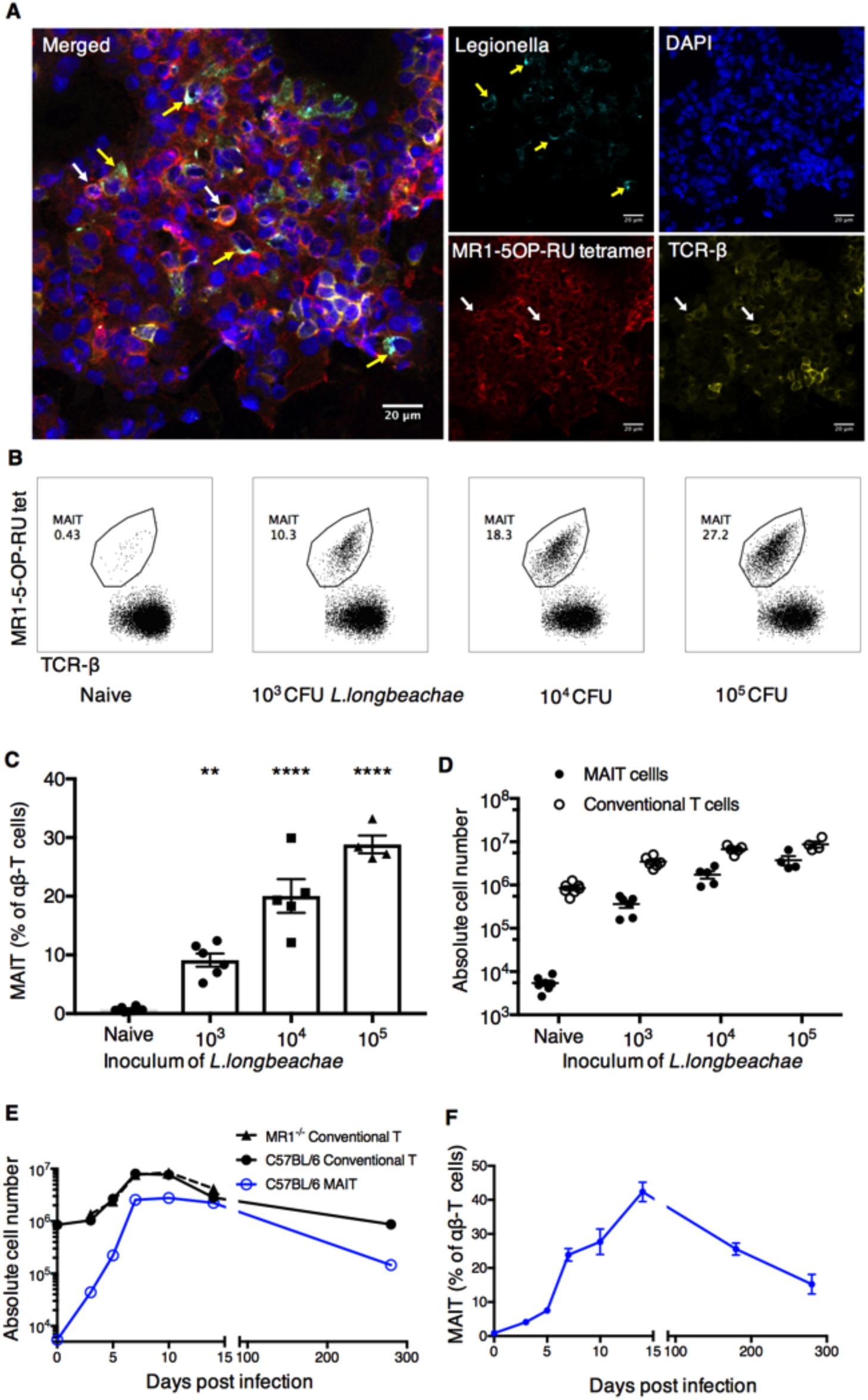
Murine pulmonary infection with *Legionella* induces long-lasting expansion of MAIT cells *in vivo* (A) Immunofluorescence micrographs of murine lungs showing TCRβ+MR1-5-OP-RU- tetramer+ MAIT cells (white arrows) adjacent to infected cells (yellow arrows) within parenchyma four days after intranasal infection with 2x10^4^ CFU *L. longbeachae* in C57BL/6 mice which had been challenged with the same inoculum 5 months previously. (B) Flow-cytometry plots showing MAIT cell percentage among TCRβ+ lymphocytes in the lungs of C57BL/6 mice either uninfected or infected with 10^3^, 10^4^ or 10^5^ CFU *L. longbeachae* for 7 days. Relative (C) and absolute (D) numbers of MAIT cells and conventional αβ T cells 7 days post infection. Absolute numbers (E), and relative percentages (F) of MR1-tetramer+ MAIT cells or conventional αβ T cells in C57BL/6 and MR1-/- mice after intranasal infection with 2x10^4^ CFU *L. longbeachae.* Experiments used 4-6 mice per group (mean±SEM) and were performed twice with similar results. B6, C57BL/6 mice; CFU, colony forming units; DAPI, 4′,6-diamidino-2-phenylindole. Statistical tests C: ** Dunnett’s P<0.01, **** P<0.0001; C all comparisons with the respective naïve groups P<0.0001 for Dunnett’s on log transformed data. See also Figures S1, S2.

Histology of lungs from mice infected with 2×10^4^ CFU of *L. longbeachae* at 7DPI demonstrated pronounced alveolar infiltration of neutrophils and macrophages, leukocytoclasia, aggregates of fibrin and accumulation of edema fluid and epithelial shedding, consistent with the typical features of human *L. pneumophila* pneumonia^15^(Supplementary Figure S2A,B). Blinded analysis of these sections using a qualitative histological score at multiple time-points post infection revealed inflammation peaked at day 7, but there were no gross histological differences in the severity of pneumonia between C57BL/6 and MR1−/− mice (Supplementary Figure S2C). To determine the cellular localization of *L. longbeachae* we measured bacterial burden in flow-sorted cells from collagenase-dispersed murine lungs 3 days post infection. Most viable bacilli localized within neutrophils, but evidence of infection of macrophages and dendritic cells was also observed (Supplementary Figure S2D).

To explore MAIT cell function we investigated the dynamics of their cytokine profile throughout infection. During acute *L. longbeachae* infection MAIT cells secreted interleukin (IL)-17A, IFN-g, GM-CSF (Figure 3A, Supplementary Figure S3) and TNF-a (data similar to INF-g, not shown). Expression of IL-17A was abundant throughout the course of the infection, whilst IFN-g secretion was significantly higher during the acute infection than in naïve cells or after resolution (each P<0.005, Figure 3B). Conversely, expression of GM-CSF was lowest during acute infection and peaked after disease resolution (P=0.0006 acute v resolution). This correlated with a shift in expression of nuclear transcription factors associated with Th1 or Th17 differentiation. In naïve mice most (81 ±4%, mean ±SD) MAIT cells expressed the orphan nuclear receptor, retinoic acid-related orphan receptor gt (RORgt) alone: a master regulator of Th17 cell differentiation (Figure 3C, 3D). A minority (13±4%) of cells expressed both RORgt and the Th1 regulator T-bet, and very few expressed T-bet alone. However, during acute infection and long-term post infection there was a marked shift in phenotype towards predominant co-expression of RORgt and T-bet in 64±5% and 69±3% of MAIT cells respectively. MAIT cells expressing T-bet alone were only observed at significant frequencies (14±3%) in acute infection. Thus the consistent secretion of IL-17A in all stages of infection and the transient increase of IFN-g secretion during acute infection reflect the changes in transcription factor profile we observed, suggesting the formation of an authentic memory pool of MAIT cells and pointing to a specific role for IFN-g in the acute response to infection.

**Figure 3.**
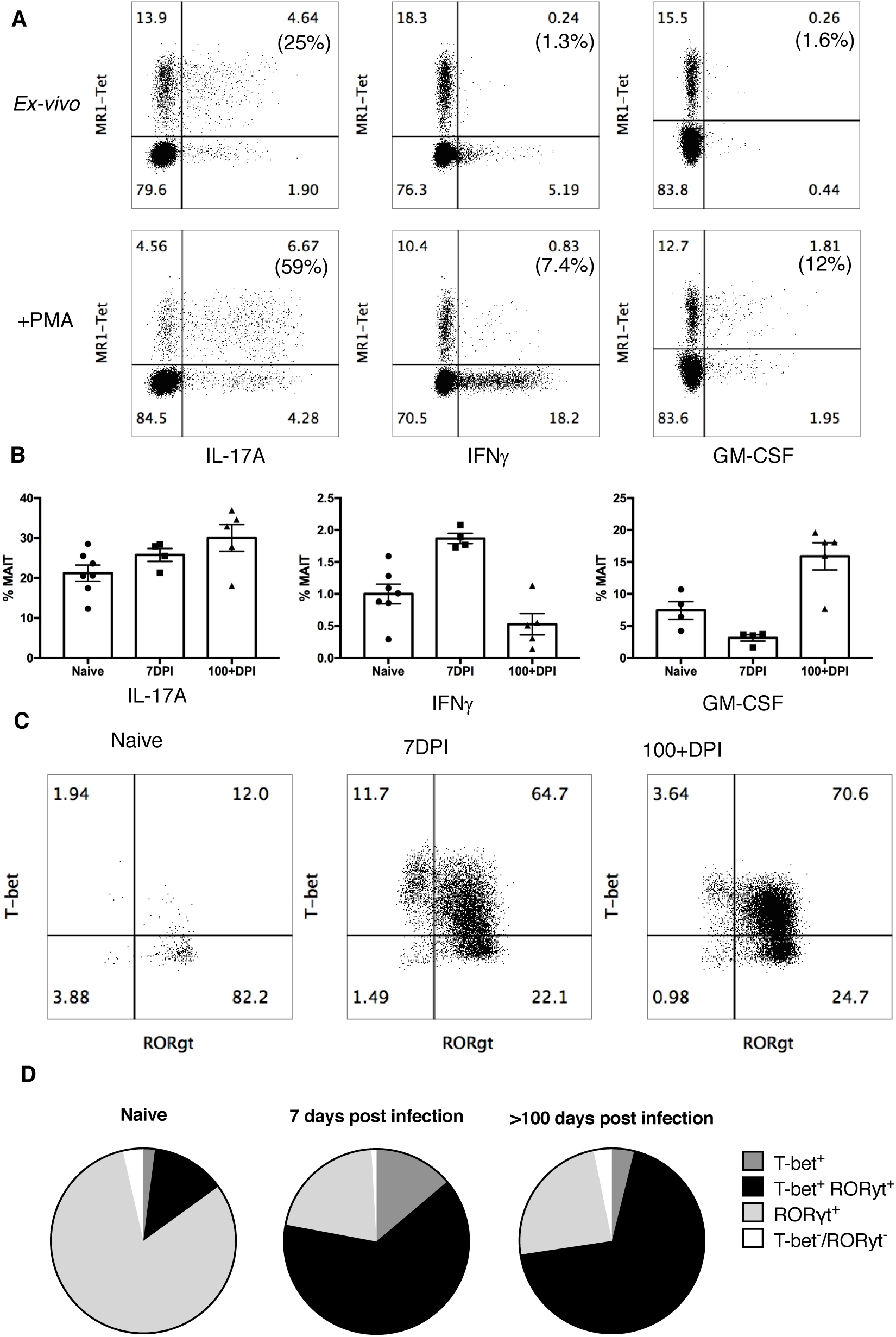
Profiles of MAIT cell cytokine expression and nuclear transcription factors vary over the course of pulmonary *Legionella* infection (A) Flow cytometry plots showing intracellular staining for IL-17A, IFN-g and GM-CSF by pulmonary TCRβ lymphocytes (non-MAIT conventional and MAIT cells) after 4h culture with or without PMA/ionomycin with bredeldin A. TCRβ+ lymphocytes were harvested from lungs of C57BL/6 mice infected with 2x10^4^ CFU *L. longbeachae* for 7 days. Percentages in brackets represent the proportion of MR1-tetramer positive MAIT cells expressing cytokine. (B) Percentages of pulmonary MAIT cells producing IL-17A, IFN-g or GM-CSF by intracellular staining, directly *ex-vivo* from C57BL/6 mice infected for 0, 7 or >100 days with 2x10^4^ CFU *L. longbeachae.* Experiments using 4-7 mice per group (mean±SEM) were performed twice with similar results. (C) Expression of T-bet and RORγt in MAIT cells from uninfected or infected C57BL/6 mice 7 or >100 days post infection (DPI). (D) Average proportion of T-bet+, double positive (DP), RORgt+ and double negative (DN) MAIT cells from uninfected or infected C57BL/6 mice at indicated date. Mean values are representative of 5-8 mice in each group. See also Figure S3.

### MAIT cell protection against life-threatening *Legionella* infection is enhanced and accelerated by prior boosting

To determine whether MAIT cells contribute to immune protection against *Legionella* we compared bacterial burden in lungs of C57BL/6 and MR1−/− mice throughout infection. Bacterial load increased by 2.5 log over the initial inoculum, peaking at 3 days postinfection (3DPI). In normal C57BL/6 mice we observed a significant difference in bacterial load but not until days 10 and 14 post infection. This was of the order of one log in CFU, consistent with relatively impaired bacterial clearance in MAIT cell deficient, MR1−/− mice (Figure 4A,B).

**Figure 4.**
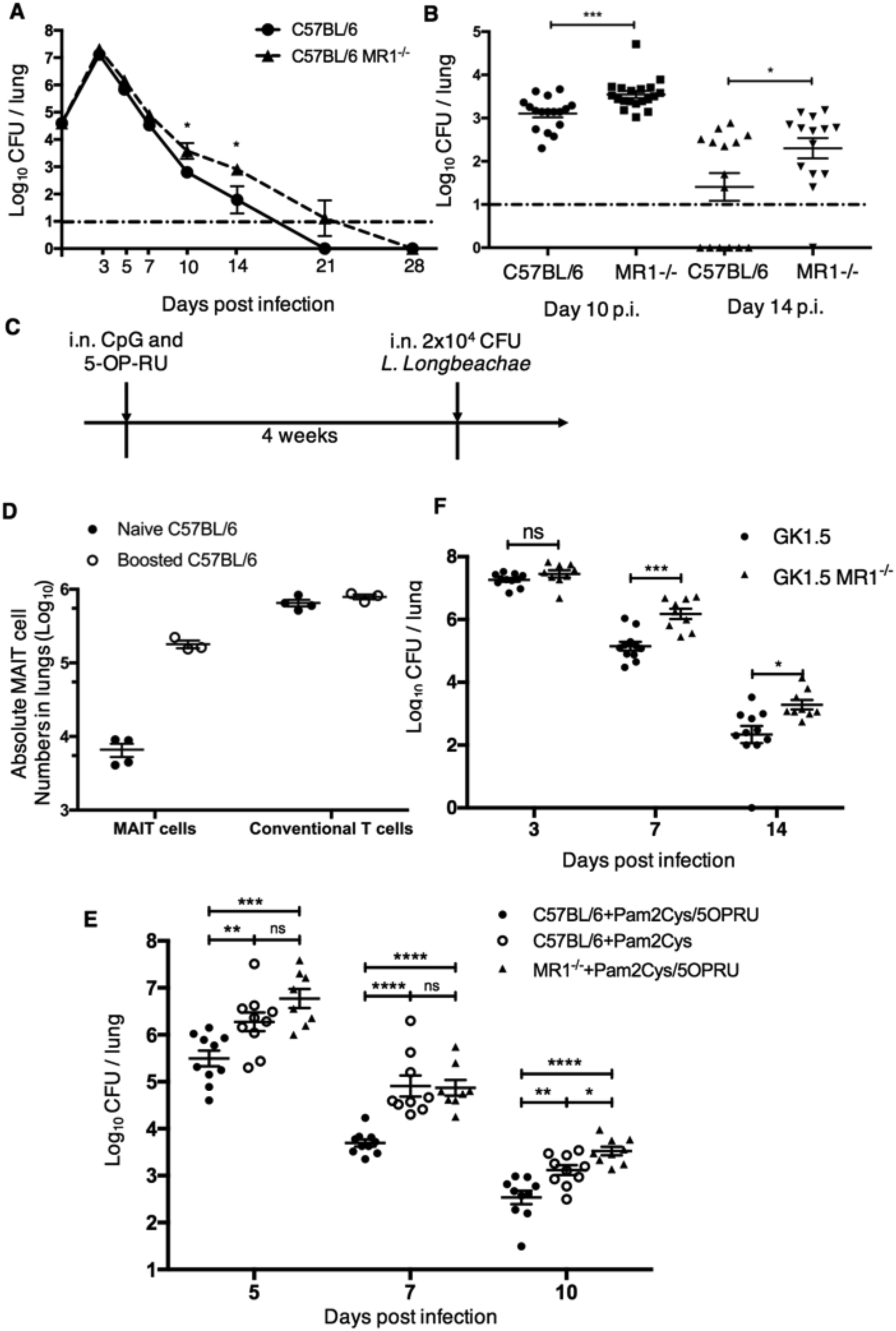
MAIT cells contribute to protection in murine *Legionella* infection in vivo, which can be accelerated by prior ligand-induced MAIT cell expansion (A,B). Bacterial load (CFU) in lungs C57BL/6 or MR1-/- mice following intranasal infection with 2x10^4^ CFU *L. longbeachae.* Dashed line represents limit of detection. (A) Bacterial load over the time-course of infection. (B). Bacterial load at days 10 and 14 post infection, from further three separate replicates. (C). Schematic for panel D and E: C57BL/6 or MR1-/- mice were treated with 20nmol S-[2,3-bis(palmitoyloxy)propyl] cysteine (Pam2Cys) and 76pmol 5-OP-RU (in 50μg intranasally 1 month before 2x10^4^ CFU *L. longbeachae* inoculation. (D) Absolute numbers of MAIT cells and conventional T cells from lungs of naïve or ligand-boosted C57BL/6 mice 30 days after administration of ligand. (E) Differences in bacterial load in lungs of C57BL/6 or ligand-boosted C57BL/6 or MR1-/- mice apparent at 5, 7 and 10 DPI. (F) Bacterial load in lungs of mice lacking CD4+ cells (GK1.5Tg) or CD4+ and MAIT cells (GK1.5Tg.MR1-/-) 3, 7 and 14 days after infection with 2x10^4^ CFU i.n. *L. longbeachae.* Pooled data (mean±SEM) from two replicates with similar results using 5-6 mice per group, compared using t tests on log-transformed data. *, P<0.05; **, P<0.01; ***, P<0.001; **** P<0.0001.

In specific pathogen-free C57BL/6 mice baseline frequencies of MAIT cells are very low^12, 13^, potentially due to lack of natural exposure to diverse environmental pathogens. We have previously shown that MAIT cells can be expanded *in vivo* by intranasal exposure to the MAIT cell ligand 5-OP-RU with a Toll-like receptor (TLR) agonist such as the TLR9 agonist CpG or TLR2 agonist S-[2,3-bis(palmitoyloxy)propyl] cysteine (Pam2Cys) to furnish a MAIT cell costimulus^13^. To understand whether MAIT cell vaccination might impact on protection observed against *Legionella* infection of the lung, we used this approach to specifically expand pulmonary MAIT cells one month prior to *Legionella* infection, without affecting conventional T cell frequencies (Figure 4C,D). Prior exposure to 5-OP-RU and CpG enhanced MAIT cell numbers in the lungs and was associated with protection against infection as reflected in a reduction in bacterial load in C57BL/6 versus MR1−/− mice (compare Figures 4A, B to Supplementary Figure S4). This protective effect became apparent earlier than observed in wild type C57BL/6 mice with reduced CFU seen on days 5 and 7 post-infection and comparable on d10 post-infection (compare Figures 4A, B to Supplementary Figure S4), as MAIT cell numbers became indistinguishable in boosted and non-boosted mice (not shown). When a direct comparison was made between MR1−/- mice, C57BL/6 mice and C57BL/6 mice that had been boosted by 5-OP-RU and Pam2Cys, bacterial burden was significantly lower on days 5, 7 and 10 post-infection in wild-type mice that had received this prior MAIT cell boosting (Figure 4E). This demonstrates the potential to augment MAIT cell-mediated protection by the prior administration of synthetic ligands as a ‘vaccine’.

These data demonstrate that MAIT cells contribute actively to *Legionella* protection in the context of an intact immune system and that this protection is more rapid and of greater magnitude when mice are first vaccinated to expand MAIT cells before infectious challenge.

### MAIT cell-mediated protection is more apparent in immune deficient mice

Studies of other intracellular pathogens have demonstrated high levels of functional redundancy in the ability of different lymphocytes subsets to control bacterial growth *in vivo^23^.* We hypothesized that by removing partially-redundant effects of other lymphocyte subsets, the protective effects of MAIT cells would become more apparent. CD4+ T cell-derived IFN-γ has been shown to play an essential role in achieving bacterial clearance of *Salmonella* Typhimurium^23^. We therefore used GK1.5 transgenic mice, which express the anti-GK1.5 antibody and are CD4+ T cell deficient^24^, and compared these with GK1.5.MR1-/- mice which lack both CD4+ cells and MAIT cells. As expected, we observed a protective effect of MAIT cells through reduced bacterial burden apparent even earlier in the course of infection than with wild type mice (statistically significant by day 7 p.i) (Figure 4F).

To further unmask the potential of MAIT cells in protection, we removed additional layers of immunity by studying the impact of adoptively transferred MAIT cells into profoundly immunodeficient Rag2^-/-^γC^-/-^ mice. We first expanded pulmonary MAIT cells by i.n. inoculation of donor mice with *S.* Typhimurium BRD509, as previously described^13^. Flow-sorted pulmonary MAIT cells from these mice were then adoptively transferred into recipient Rag2^-/-^γC^-/-^ mice in which Rag2 and the common *g* chain are deleted, leading to absence of T, B and NK cells. After transfer, administration of anti- CD4 and anti-CD8 mAbs was used to further deplete any residual contaminating conventional T cells (Figure 5A). After adoptive transfer, MAIT cells expanded spontaneously to generate a stable population by two weeks (Figure 5B, Supplementary Figure S5A) which expressed the nuclear transcription factor and master regulator of innate-like T cell development promyelocytic leukemia zinc finger(PLZF) (Figure S5B) ^25^

**Figure 5.**
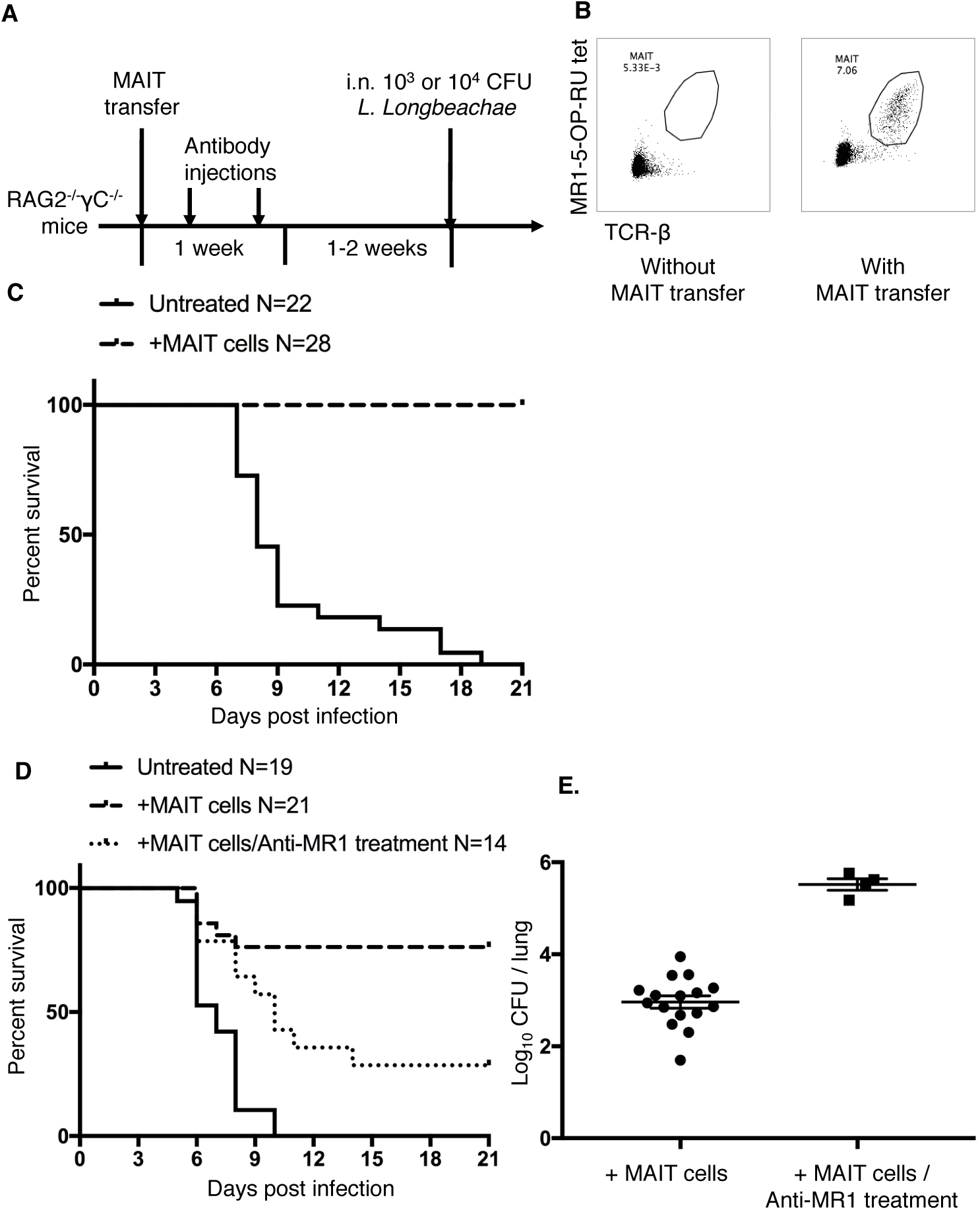
Adoptive transfer of MAIT cells rescues Rag2-/-yc-/- mice from fatal pulmonary *Legionella* infection (A) Schematic of protocol: 10^5^ pulmonary MAIT cells from C57BL/6 mice previously infected with 10^6^ CFU *S.* Typhimurium BRD509 for 7 days to expand the MAIT cell population were sorted and transferred intravenously into Rag2-/-gC-/- mice, followed by intraperitoneal anti-CD4 and anti-CD8 antibody injection (0.1mg each) twice within 1 week to deplete contaminating conventional T cells. After 2 weeks, mice were infected with 10^3^ or 10^4^ CFU i.n. of *L. longbeachae.* (B) Representative plots showing live (7AAD-) hematopoietic (CD45.2+) cells with percentages of MAIT cells in the lungs of Rag2-/-gC-/- mice which were untreated or were recipients of adoptively-transferred MAIT cells. (C) Survival of *Legionella*-infected untreated or MAIT cell-recipient Rag2- /-gC-/- mice after 10^3^ CFU i.n. *L. longbeachae* infection. (D) Survival of Legionella- infected untreated or MAIT cell-recipient Rag2-/-gC-/- mice after 10^4^ CFU i.n. *L. longbeachae*, with or without MR1 blockade. One group received 0.25mg anti-MR1 monoclonal antibody alternate days after infection. (E) Pulmonary bacterial load in surviving Rag2-/-gC-/- mice in (D) 23 DPI. Pooled data (mean±SEM) from two replicates with similar results, each with 7-12 mice per group. See also Figure S5.

Strikingly, the presence of adoptively-transferred MAIT cells was sufficient to rescue completely Rag2-/-γC-/- mice from fatal infection with 10^3^ CFU *L. longbeachae* (Figure 5C, X_2_ P<0.0001) in the absence of other components of adaptive immunity. Using a higher inoculum (10^4^ CFU) we observed this protection was reduced by blockade with anti-MR1 mAb, which was associated with significantly reduced survival (X_2_ P=0.005) and with increased bacterial load amongst surviving mice (P=0.0004), consistent with an MR1-dependent mechanism (Figure 5D,E).

### MAIT cell-protection is dependent on IFN-g

To determine the mechanism by which MAIT cells provide this protection we used adoptive transfer of MAIT cells from mice with deficiencies in cytotoxic capability or in pro-inflammatory cytokines. The protective effect of MAIT cells on both survival of Rag2^−/−^γC^−/−^ mice or on bacterial burden was not impaired in MAIT cells lacking the cytolytic proteins perforin or granzymes A and B, nor in MAIT cells unable to express IL-17A or TNF-a (Figure 6A,B). We observed a small increase in bacterial burden when transferred MAIT cells were deficient in GM-CSF (0.49 log-fold difference in CFU, P=0.026), but this was not associated with significant differences in survival. By contrast protection was critically dependent on MAIT cell derived IFN-g, with decreased survival (P<0.0001) and a 2.8 log-fold increased bacterial burden (P<0.001) when MAIT cells were deficient in IFN-g. Furthermore these mice all succumbed to *Legionella* infection by day 37 p.i..

**Figure 6.**
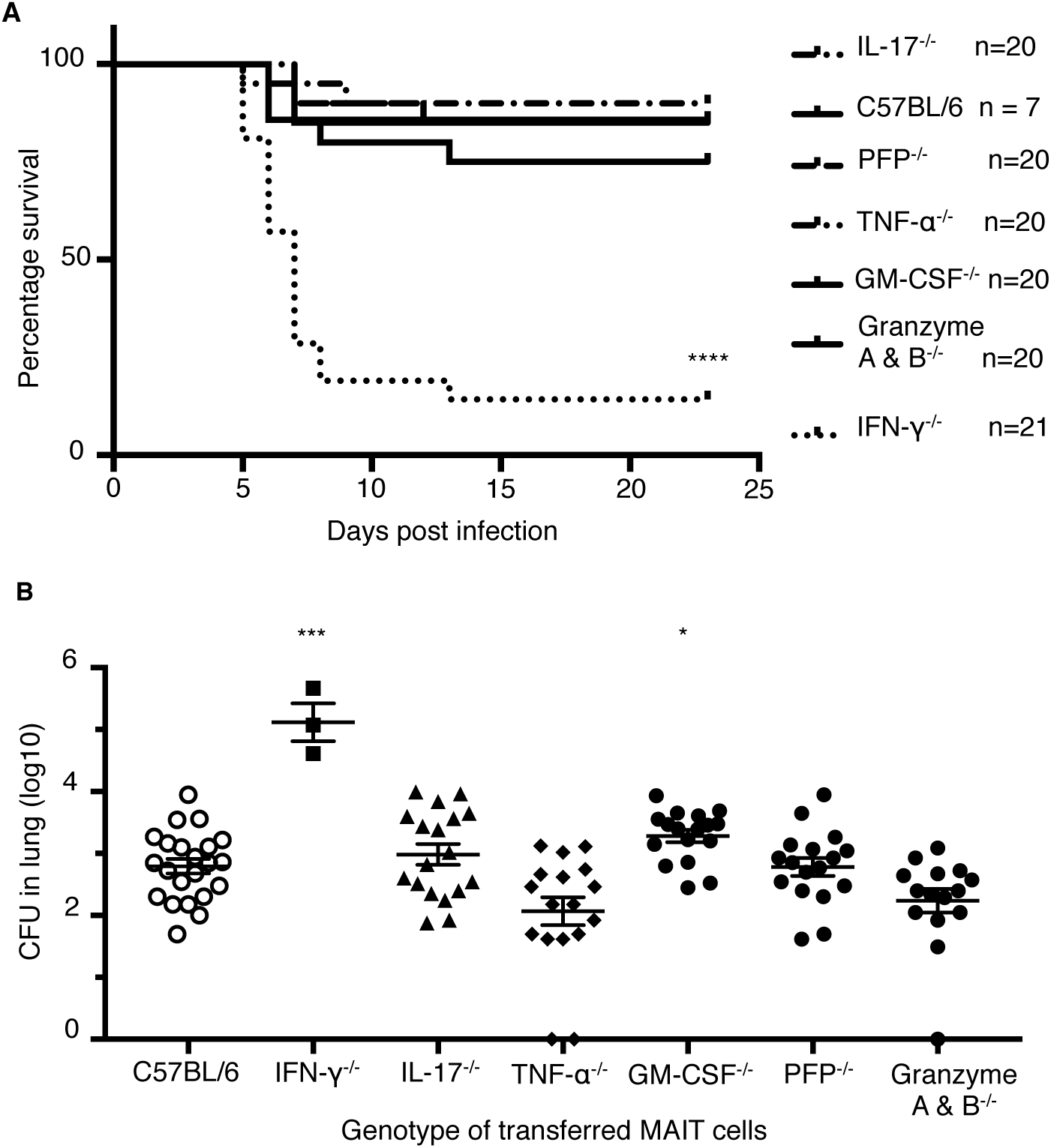
Protection of Rag2-/-gc-/- mice from fatal pulmonary *Legionella* infection is dependent on IFN-γ (A) Survival of Rag2-/-gC-/- mice adoptively transferred with pulmonary MAIT cells generated in different mouse strains after 10^4^ CFU i.n. *L. longbeachae.* ****, Gehan- Breslow-Wilcoxon P<0.0001. (B) Pulmonary bacterial load in surviving Rag2-/-gC-/- mice in (A) 23 DPI. Pooled data (mean±SEM) from two replicates with similar results, each with 7-13 mice per group. Fig B shows mean. PFP, perforin. Bonferroni-corrected t- tests *, P<0.05; **, P<0.01; ***, P<0.001.

The use of adoptive transfer of *in vivo* expanded MAIT cells provides compelling evidence that MAIT cells can confer protection against important human pathogens and demonstrates this protection depends upon their capacity to produce IFN-γ and to a lesser extent GM-CSF.

## Discussion

Our findings show that MAIT cells are activated and proliferate in response to *Legionella* infection, leading to enhanced immune protection *in vivo* that is dependent on IFN-γ and GM-CSF. This protection is evident earlier and of greater magnitude if mice are first vaccinated to expand and prime MAIT cells which are otherwise present in small numbers in normal mice. Protection by MAIT cells is characterized by more rapid reduction in bacterial loads and is MR1-dependent suggesting mediation via antigen- specific activation. Remarkably, MAIT cell protection against *Legionella* was nonredundant and even evident in fully immune competent mice. The protective effect of MAIT cell immunity became more evident as layers of immunity were removed in host mice, firstly in GK1.5 mice lacking only CD4+ T cells and then in more profoundly immundeficient Rag ^−/−^γC^−/−^ mice, lacking conventional T cells, B cells and NK cells. This observation is important given that studies of primary immunodeficiencies^26^ imply redundancy of different lymphocyte subsets is a typical feature of pathogen immunity especially for innate mechanisms such as NK cells and innate lymphoid cells. Indeed, in the absence of B, T and NK cells MAIT cells were absolutely critical for survival in *Legionella*-infected mice revealing their important potential in compromised hosts. As this mechanism was dependent on MR1, which presents small molecules derived from riboflavin biosynthesis^3-5^, this demonstrates *in vivo* the potential for control of *Legionella* by detection of riboflavin metabolites.

These observations suggest how the contribution of MAIT cells to immune protection may be critical to survival in clinical, naturally-occurring severe infection. The mortality we observe from *Legionella* does not coincide with the time of peak bacterial load - on day 3 post-infection - but later, between days 6 and 14 post-infection, at which point MR1-/- mice had 0.78 to 1.1-log fold higher bacterial load than wild-type mice. In essence, MAIT cells may be the difference between life and death in knife-edge infections where host immunity is partially compromised by comorbidities or predisposing factors, or where patients are exposed to large bacterial doses.

Although MAIT cells have both cytotoxic activity^27^ and the ability to rapidly produce pro-inflammatory cytokines including interleukin IL-17A, TNF-a and IFN-g^10^, the protective effect of MAIT cells against *Legionella* infection was not dependent on TNF-a or IL-17A, but instead relied upon the capacity of MAIT cells to secrete IFN-g and GM- CSF. This is consistent with a study of *Francisella* infection^28^ where GM-CSF reduces bacterial burden late in the course of infection, although this did not translate into a significant survival difference. Nor did the protective effect depend on the key cytotoxic effector molecules: perforin and granzymes A/B. This is in contrast to work suggesting MAIT cell cytotoxicity is important for control of *Shigella-infected* HeLa cell lines *in vitro^27^*.

As *Legionella* infects inflammatory cells, particularly neutrophils and macrophages, rather than epithelia, the essential immune function required of MAIT cells in our system is likely the IFN-g-stimulated enhancement of bactericidal activity within these cells in which phagosome function has been subverted. Our findings of IFN-g production upon MAIT cell activation are consistent with other reports^6,10,14,27^, and accord with reports of a role for MAIT cell-derived IFN-g in limiting growth of *Francisella tularensis* in bone marrow-derived macrophages *in vitro^14^.* IFN-g has also been shown to enhance bactericidal activity of neutrophils via multiple mechanisms including enhancement of oxidative burst, nitric oxide production, antigen presentation, phagocytosis, and upregulation of CD80/86 co-stimulation and T cell-recruiting cytokines and chemokines^29^. Moreover, given that IFN-g is critical also for protection against mycobacterial disease including *M. tuberculosis (M.tb)*, it is likely that this early production of MAIT cell-derived IFN-g may be an important and non-redundant component of protection against mycobacteria. Indeed *in vitro* MAIT cell-derived IFN-g inhibits growth of *Bacillus Calmette-Guerin* (BCG) in macrophages^30^, and the MR1- MAIT cell axis has been linked to susceptibility to BCG in mice^30^ and to *M.tb* in humans^31^ and mice^32^.

A striking feature of MAIT cell biology is the very low frequencies of MAIT cells we observe in blood or lungs in naïve mice^12, 13^, in contrast to the marked and long-lived expansion induced by a single infection in our model. Although antigen-naïve MAIT cells have some intrinsic effector capacity^33^, the delay between initial microbial exposure and peak MAIT cell frequency may be critical in providing a window of opportunity for a pathogen to exploit^14^. This notion is consistent with our observation that MAIT cell protection can be accelerated by prior expansion of the pulmonary MAIT cell population using intranasal synthetic 5-OP-RU and an appropriate TLR agonist. Notably, MAIT cell frequencies are low in early childhood^34^, suggesting the potential to enhance the immunogenicity of vaccines given in early life by incorporating such MAIT cell ligands in combination with TLR stimulation, which might for instance promote the recruitment of inflammatory monocyte differentiation via MAIT cell-derived GM-CSF^28^. A protective effect of expanding MAIT cells could contribute to heterologous protection afforded by neonatal BCG vaccination against other, unrelated classes of pathogens^35^. Vaccination with MAIT cell ligands might help resolve chronic infections where MAIT cell frequencies may be reduced due to other therapies^11^, comorbidities or activation induced cell death^36^.

Our findings define a significant role for MAIT cells in pulmonary host defense against a major human pathogen. We reveal layers of immunological redundancy likely to mask the contribution of MAIT cells in many situations of infectious challenge, but suggest a critical role for MAIT cells becomes apparent in a crisis situation - as reflected here in a high infecting inoculum, or in the face of compromised specific immunity - in which the gulf between survival and death is finely balanced. Moreover, we demonstrate the mechanism of this MAIT cell protection is IFN-γ dependent and enhanced by GM-CSF. Due to the pleiotropic roles of IFN-γ and the conservation of the riboflavin pathway across many species, this mechanism is likely broadly effective against other major human intracellular pathogens, and may prove as relevant to the later stages of infection as to the initial, acute phase characterized by innate immune responses countering rapid pathogen replication. We have shown this immunity can be augmented by exposure to MR1 ligands suggesting this mechanism might have potential for preventive or therapeutic benefit.

## Materials and Methods

### *In vitro* activation assays

Jurkat cells expressing a MAIT TCR comprising the TRAV1-2-TRAJ33 a-chain and TRBV6-4 β-chain (Jurkat.MAIT) were co-incubated at 10^5^ cells per well in 96-well U-bottom plates with an equal number of class I reduced (C1R) antigen presenting cells (APCs) expressing MR1 (C1R.MR1)^3^ for 16 h in RPMI-1640 media (Gibco) in supplement and 10% foetal calf serum (FCS)) (RF10) media at 37°C, 5%CO_2_. Cells were stimulated for 16 h with bacterial lystates prepared using repeated ultrasonication of bacteria in log-phase growth. Cells were stained with anti-CD3-APC and anti-CD69-PE Abs and 7AAD before flow cytometric analysis. Activation of Jurkat.MAIT cells was measured by an increase in surface CD69 expression.

For human MAIT cell assays, peripheral blood mononuclear cells (PBMCs) were stained with anti-CD3-PEAF594, CD161-PECy7, TCR Va7.2 (TRAV1-2)-APC. Va7.2+ cells were enriched with anti-APC magnetic beads and the CD3+Va7.2+CD161+ population were isolated by flow cytometry and cultured at 10^4^ cells/well for 16 or 36 hours in penicillin-free media containing streptomycin and gentamicin with 5x10^4^ THP1 cells which had been first infected for 3 hours (CD69 assays) or 27 hours (intracellular cytokine staining) with different multiplicities of infection (MOI) of live *L. longbeachae* NSW150. Control wells contained 5-OP-RU 10nM or *L. longbeachae* NSW150 at MOI 100 which had been heat killed for 10 minutes at 67°C. CD69 upregulation was measured by surface staining for CD69-APCCy7 and 5-OP-RU loaded human MR1 tetramer-PE. For intracellular cytokine expression cells brefeldin A was added for the last 16 hours, cells fixed, permeabilized (using BD Fixation/Permeabilization Kit (BD, Franklin Lakes, NJ) and stained with MR1 tetramer, CD3-PEAF594, Zombie yellow, IFNg-FITC, and TNFa-Pacific blue.

### Generation of THP1:MR1- cell line

THP1:MR1- cell lines were generated by targeted deletion of MR1 using LentiviralCRISPRv2 which was a gift from Feng Zhang (Addgene plasmid # 52961)^37^. The plasmid was digested with *BsmB1* (Fermentas), dephosphorylated and purified using gel electrophoresis and the Ultraclean DNA isolation kit (MO Bio laboratories, Carlsbad CA). Short guide RNAs (ACCTCTCATCATTGTGTTAA) were ligated and the reaction product used to transform Stbl3 *E.Coli.* DNA was purified from transformed colonies (QIAprep spin miniprep) and used to transfect HEK293T cells with LentiviralCRISPRv2 vector and packaging vectors. Supernatants were used to transduce parent THP1 cells and cells were selected using puromycin resistance and single cell sorting using anti-MR1 antibody (8F2.F9) after upregulation of MR1 using acetyl-6FP. MR1 knockouts were then verified using surface staining and western blotting.

### Immunofluorescence microscopy

8 μm sections of cryopreserved, unfixed lung tissue were submerged into ice-cold acetone for 10 min, air dried and then re-hydrated in PBS for 10 min. Endogenous biotin block was performed using Biotin/Avidin blocking kit (Thermo Fisher, Waltham MA) according to the manufacturer’s instructions. Serum-free protein block (DAKO, Carpinteria, CA) was applied for 15 min, followed by 30 min blocking with 10% normal donkey serum. Sections were subsequently blocked with Murine MR1-6FP tetramer (Nil PE) for 1 hour at room temperature. Murine MR1-5-OP-RU tetramer-PE (25 μg/ml in 2% bovine serum albumin/PBS) was applied for 1 h at room temperature in the dark, sections washed with PBS, fixed with 1% paraformaldehyde for 10 min, washed again and stained with a cocktail containing polyclonal rabbit anti-Legionella antibody (gift from Dr Hayley Newton, Department of Microbiology and Immunology, University of Melbourne) and Goat anti PE (KPL). After 1 hr, sections were washed and stained with Donkey anti Goat-AlexaFluor 568 (Life Technologies), Donkey-anti-Rabbit-DyLight 680 (Life Technologies) and Alexa Fluor 647-conjugated Rat anti mouse TCR-β (Biolegend). Nuclei were counterstained with Hoechst 33342 (Life Technologies). Sections were mounted with Prolong Gold mounting medium (Life Technologies). Image acquisition was performed on Zeiss LSM 710 confocal microscope using Zen software (Zeiss, Oberkochen, Germany) The resultant images were further analysed using FIJI Image J software^38^ (UW-Madison).

### Animal models

Mice were bred and housed in the Biological Research Facility of the Peter Doherty Institute (Melbourne, Victoria, Australia). MR1-/- mice were generated by breeding Va19iCa-/-MR1-/- mice^39^ (from Susan Gilfillan, Washington University, St Louis School of Medicine, St Louis, MO) with C57BL/6 mice and inter-crossing of F1 mice. The genotype was determined by tail DNA PCR at the MR1 locus as previously described^13^. Granzyme A/B-/- and Perforin-/- mice were purchased from Joe Trapani (Victorian Comprehensive Cancer Centre, Melbourne). GK1.5 mice were crossed onto the MR1-/- background to generate GK1.5.MR1-/- mice which lack CD4+ cells and MAIT cells. Male mice aged 6-12 weeks were used in experiments, after approval by the University of Melbourne Animal Ethics Committee (1513661).

### Intranasal infection

Intranasal (i.n.) inoculation with *L. longbeachae* or antigens (76 pmol 5-OP-RU) and TLR agonist (either 20 mg CpG or 20nmol Pam2Cys) in 50 μ! per nares was performed on isofluorane-anesthetized mice. For blocking experiments, mice were given 250 μg anti-MR1 (26.5 or 8F2.F9)^21, 40^ or isotype control mAbs in 200 μl PBS, once (i.v or intraperitoneally) 1 day prior to inoculation and three times (d1, d3, d5) post inoculation. Mice were killed by CO_2_ asphyxia, the heart perfused with 10ml cold RPMI and lungs were taken.

To prepare single-cell suspensions lungs were finely chopped with a scalpel blade and treated with 3mg/ml-1collagenase III (Worthington, Lakewood, NJ), 5 μg/ml DNAse, and 2% fetal calf serum in RPMI for 90 min at 37°C with gentle shaking. Cells were then filtered (70μm) and washed with PBS/2% foetal calf serum. Red blood cells were lysed with hypotonic buffer TAC (Tris-based amino chloride) for 5 min at 37°C. Approximately 1.5x10^6^ cells were filtered (40μm) and used for flow cytometric analysis.

### Determination of bacterial counts in infected lungs

Bacterial colonization was determined by counting colony-forming units (CFU) obtained from plating homogenized lungs in duplicate from infected mice (x5 per group) on buffered charcoal yeast extract agar containing 30μg/ml streptomycin and colonies counted after 4 days at 37°C under aerobic conditions.

### Adoptive transfer

As MAIT cell frequencies are low in naïve C57BL/6 mice, prior to adoptive transfer experiments MAIT cell populations were expanded by intranasal infection with 10^6^ CFU *S.*Typhimurium BRD509 in 50μ! PBS for 7 days as previously described^13^. After 7 days, mice were sacrificed, single cell suspensions prepared and live CD3+CD45+MR1-5-OP-RU tetramer+ cells sorted using a BD FACS Aria III. 10^5^ MAIT cells were injected into the tail veins of recipient mice which then received 0.1 mg each of anti-CD4 (Gk1.5) and anti-CD8 (53.762) mAb i.p on days 2 and 5 or 6 to control residual conventional T cells. Mice were rested for 2 weeks post transfer to allow full expansion of the MAIT cell population prior to subsequent infectious challenge. Mice were weighed daily and assessed for visual signs of clinical disease, including inactivity, ruffled fur, labored breathing, and huddling behavior. Animals that had lost ≥20% of their original body weight and/or displayed evidence of pneumonia were euthanized.

### Statistical analysis

Statistical tests were performed using the Prism GraphPad software (version 7.0 La Jolla, CA). Comparisons between groups were performed using Student’s t-tests or Mann-Whitney tests as appropriate unless otherwise stated. Survival curves were compared using the Gehan-Breslow-Wilcoxon method for multiple groups. Flow cytometric data analysis was performed with FlowJo10 software (Ashland, OR).

### Reagents

Human peripheral blood mononuclear cells (PBMC) were obtained from the Australian Red Cross Blood Service (ARCBS) (University of Melbourne Human Research Ethics Committee 1239046.2). Healthy human lung explant tissue was obtained via the Alfred Lung Biobank program and ARCBS from organs not suitable for donation (Blood Service HREC 2014#14 and University of Melbourne Human Research Ethics Committee 1545566.1).

### Compounds, immunogens and tetramers

5-OP-RU was prepared as described previously^18^. CpG1688 (Sequence: T*c*C*A*T*G*A*C*G*T*T*C*C*T*G*A*T*G*C*T (*phosphorothioate linkage) nonmethylated cytosine-guanosine oligonucleotides was purchased from Geneworks (Thebarton, Australia). Murine and human MR1 and β2-Microglobulin genes were expressed in *Escherichia coli* inclusion bodies, refolded, and purified as described previously^41^. MR1-5-OP-RU tetramers were generated as described previously^4^.

### Bacterial strains

Cultures of *Legionella pneumophila* JR32 and *Legionella longbeachae* NSW150 were grown at 37°C in buffered yeast extract (BYE) broth supplemented with 30- 50μg/ml streptomycin for 16 hour to log-phase (OD600 0.2-0.6) with shaking at 180 rpm. For the infecting inoculum, bacteria were re-inoculated in prewarmed medium for a further 2-4 h culture (OD_600_ 0.2-0.6) with the estimation that 1 OD_600_=5x10^8^/ml, sufficient bacteria were washed and diluted in phosphate buffered saline (PBS) with 2% BYE for i.n. delivery to mice. A sample of inoculum was plated onto BYCE with streptomycin for verification of bacterial concentration by counting colony-forming units.

For infection of adoptive transfer donor-mice with *Salmonella* Typhimurium BRD509 cultures were prepared as previously described^13^.

### Antibodies and flow cytometry

Antibodies against murine CD4 (GK1.5, APC-Cy7), CD19 (1D3, PerCP-Cy 5.5), CD45.2 (104, FITC), IFNg (XMG1.2, PE-Cy7), Ly6G (IA8, PECy7), TCRβ (H57-597, APC or PE), TNF-a (MP6-XT22, PE), GM-CSF (MP1-22E9, PE) and IL-17A (TC11- 18H10, PE) were purchased from BD (Franklin Lakes, NJ). Antibodies against CD8a (53-6.7, PE), PLZF (Mags.21F7, PE), RORgt (B2D, APC), T-bet (4B10, PE-Cy7) and MHCII (M5/114.15.2, AF700) were purchased from eBioscience (San Diego, CA). Abs against CD19 (6D5, BV510), F4/80 (BM8, APC), CD11b (M1/70, FITC), CD11c (N418, BV786), CD31 (PCAM, MEC13.3, PerCPCy5.5), CD62L (Mel-14, FITC), CD64 (X54- 5/71, BV711), CD146 (ME-9F1, PerCPCy5.5), CD326 (G8.8, EpCAM, APC-Cy7) were purchased from Biolegend (San Diego, CA). Blocking Ab (26.5, 8F2.F9) and isotype controls (3E12, 8A5) were prepared in house. To block non-specific staining, cells were incubated with MR1-6FP tetramer and anti-Fc receptor (2.4G2) for 15 min at room temperature and then incubated at room temperature with Ab/tetramer cocktails in PBS/2% fetal calf serum. 7-aminoactinomycin D (5 μ! per sample) was added for the last 10 min.

Antibodies against human CD3 (UCHT1, PE-AlexaFluor594), TCR-Va7.2 (3C10, APC), CD161 (HP-3G10, PE-Cy7), TNF-α (Mab11, Pacific Blue), and viability dye (Zombie Yellow) were purchased from Biolegend. Antibodies against IFN (25725.11, FITC) and CD69 (FN50, PE) were purchased from BD, and anti-CD3 (UCHT1, APC) from eBioscience.

Cells were fixed with 1% paraformaldehyde prior to analysis on LSRII or LSR Fortessa or Canto II (BD Biosciences) flow cytometers. For intracellular cytokine staining Golgi plug (BD Biosciences) was used during all processing steps. Cells stimulated with PMA (phorbol 12-myristate 13-acetate;)/ionomycin (20 ng/ml, 1μg/ml, respectively) for 3 h at 37°C were included as positive controls. Surface staining was performed at 37°C, and cells were stained for intracellular cytokines using the BD Fixation/Permeabilization Kit (BD, Franklin Lakes, NJ) or transcription factors using the transcription buffer staining set (eBioscience) according to the manufacturers’ instructions.

## Acknowledgments

This research was supported by National Health and Medical Research Council of Australia (NHMRC) Program Grants 1113293, 1016629 and 606788, a Project Grant 1120467 as well as a Merieux Research Grant. T.S.C.H. was supported by a Wellcome Trust Postdoctoral Research Fellowship (104553/z/14/z). A.J.C. was supported by an ARC Future Fellowship. S.B.G.E. was supported by an ARC DECRA Fellowship. J.R. was supported by an NHMRC Australia Fellowship, D.P.F. was supported by an NHMRC Senior Principal Research Fellowship. H.W. is supported by a Melbourne International Engagement Award (University of Melbourne). C.D’S. is supported by a Melbourne International Research Scholarship and a Melbourne International Fee Remission Scholarship (University of Melbourne). We thank Dr Wei-Jen Chua and Prof Ted Hansen for their kind provision of 8F2.F9 and 26.5 mAbs, and Prof David Jackson for Pam2Cys. We are grateful to Prof Francis Carbone for critical review of the manuscript.

## Author contributions

HW, CDS, TSCH, XYL, LK, TLP, SBG, BSM, ZC, AWS performed the experiments and analyzed the data. ZC, TSCH, HW, JM, AC, RS designed the experiments and managed the study. NW, DPF, YI, JG, GW, LK-N, JR, LL, JYWM provided essential reagents and intellectual input. TSCH, AC, ZC, JM conceived the work and wrote the manuscript which was revised and approved by all authors.

### Materials and Correspondance

Correspondence and material requests should be addressed to J McCluskey.

## Supplementary Information

Supplementary Figures S1-S5

